# Automated sleep scoring in hibernating and non-hibernating American black bears

**DOI:** 10.1101/2025.03.31.646262

**Authors:** Øivind Tøien, Elsa Cecile Pittaras, Yi-Ge Huang, Paul J. N. Brodersen, Giancarlo Allocca, Brian M. Barnes, H. Craig Heller

## Abstract

Hibernating bears show remarkable metabolic suppression. Their decline in core body temperature (T_b_) is moderate (from 38°C to 30-35°C), but their metabolism declines as much as 75%. To understand the role of sleep in this hypometabolic state, we recorded biotelemetrically EEG, EOG and EMG data over 3500 days from 16 captive American black bears in and out of hibernation under semi-natural conditions. This data set is too large to score manually for Wake, REM- and NREM sleep, so we tested two machine learning classifiers: (1) Somnotate trained on multiple one-day recordings, and (2) Somnivore, trained on a small subset from each recording. As automated scoring methods have not been applied to hibernating species before, a major concern is the effect changing brain temperature has on the EEG and on the machine learning based detection. Therefore, we selected reference data using consensus by 3 manual sleep scorers from each of 6 bears, two one-day recordings at the highest and lowest body temperatures during hibernation when T_b_ was oscillating in multiday cycles, and a non-hibernating one-day recording in summer. Somnotate results were excellent when trained separately for hibernating and non-hibernating data. Training Somnotate separately for high and low T_b_ within hibernation did not improve results further. Sleep times in hibernation were about 2x that in summer for both automated scores and manual scores (p<0.0001). There were no significant differences in occupancy of vigilance states between automated and manual scores in hibernation (p>0.05), but a small overestimate of sleep time in summer (p<0.05). Both applications yielded F-measures against manual scores in the 0.90-0.98 range. Outliers in the 0.67-0.88 range were correlated between the two applications, indicating that specific files are more challenging to annotate. We conclude that both applications have accuracies approaching that of manual scorers when trained on high quality data.

## Introduction

Bears are an unexplored and interesting model for studying therapeutic hypometabolism and the role of sleep in that adaptation. In the hypometabolic state of hibernation black bears have been found to suppress metabolic rate to 25% of normal basal rates, while their core body temperatures (T_b_) on average decrease only 5.5°C during hibernation. But, their T_b_s do not stay at a stable level, but vary in multiday body temperature cycles in the range of 30-35°C [1, 2]. Animal studies typically classify sleep into stages of non-REM sleep (NREM), rapid eye movement sleep (REM) and Wake [3], with NREM characterized by domination of medium to large amplitude delta waves caused by synchronization of neuronal firing across cortex, REM dominated by low amplitude alpha waves and presence of bipolar EOG spikes and lack of EMG activity, and Wake with irregular high-frequency, low-amplitude EEG patterns and random EMG activity. Studies in marine mammals have also defined a stage of Drowsiness [4, 5]. Human studies additionally classify NREM further into sleep stages N1, N2 and N3 according to the American Academy for Sleep Medicine (AASM) standards [6, 7]. With the exception of certain hibernating dwarf lemur species [8], EEG based sleep studies have previously only been published for smaller hibernators that go into torpor through a period of sleep but remain at too low temperatures during deep torpor to record brain activity that can be used for classification of sleep stages. Thus sleep stages can typically only be classified in these hibernators during the episodic arousal episodes when animals return to euthermic levels of T_b_ for one or two days [9, 10, 11, 12]. While functionality of the increased deltawave at early arousal caused much controversy among these studies as to its function or even if it can be classified as sleep, in contrast, bears in hibernation do not show torpor-arousal cycles and remain at high enough body temperatures to be responsive to disturbance at any time, and because their brain temperatures do not drop below the low 30 °C [1, 2], their vigilance states can be scored throughout hibernation.

Presumed without data, bears are often said to be “asleep” during hibernation, but in this context it refers to the hibernating state (“winter dormancy”). No sleep studies based on brain wave activity existed in a large hibernator before we started collecting a very extensive (>3500 days) repository of polysomnographic data in American black bears (*Ursus americanus*) kept in outdoor enclosures in Fairbanks, Alaska. Using telemetry implants we recorded global cortical EEG, EOG, EMG, ECG, T_b_, breathing and respirometry (while dens were closed) and in some of the recordings blood pressure, using telemetry implants. In contrast to small hibernators, there is need to analyze sleep architecture though the whole 5-6 month hibernation period and transitions in and out of the hibernation season to understand role of sleep in the hypometabolism of hibernating bears. Through the years of our study, our polysomnographic recordings produced too much data to score manually, and at the time data were collected reliable methods to automatically score these data in their entirety were not available. A broad range of machine learning techniques have become available for sleep classification more recently, and a number of them have been reviewed [13, 14, 15, 16]. The majority of the studies aimed at verifying machine learning based sleep classification have been performed on human data and standard laboratory animals, e.g. mice and rats. Previous studies have not verified machine learning based sleep classification of EEG recordings from hibernating animals. Thus, before any automatic scoring technique is applied to analyze these data on a larger scale, the integrity of the analysis must be verified. The variations in T_b_ of hibernators has the potential to affect the frequency distribution of the EEG signals [17], and since machine learning techniques are typically based on a first step of convolution to extract the frequency distributions of the signals [14], there could be potential problems with correct detection if the machine learning routines are trained on data recorded at different T_b_‘s than occurred during the analyzed recording. For the present study we selected two different machine learning based automatic sleep qualifiers based on their previous successful verifications on animal data: (1) open source Somnotate, a probabilistic classifier based on a combination of linear discriminant analysis with a hidden Markov model that has previously been tested on mouse data [18], and (2) Somnivore, which applies proprietary assisted machine learning algorithms that have been successfully tested on a wide range of human data and animal data from mice, rats and pigeons [19]. The purpose of the present study is to evaluate the integrity of automated sleep scoring using these two applications on data from American black bears (*Ursus americanus*) in and out of hibernation.

Due to the dissimilarity in how the two programs were designed, it was not feasible and we did not intend to perform a comparison between the two qualifiers on terms of similar training and testing conditions. While both programs can perform batch scoring on multiple files after training, Somnivore was primarily designed to be trained on a minimal amount of data from a one day recording to be tested (assisted machine learning) and currently cannot easily be trained on data from multiple files. Somnotate is able to accumulate training data from an unlimited number of files and multiple animals to generate machine learning models that provide robustness against incursion of incorrect data in the training data set and can be applied across multiple animals. For testing integrity and robustness of training models, Somnotate can automate use of holdout testing so that each file to be tested is excluded from the training data [18] and the results can then be compared to the non-holdout training model. Holdout testing would typically not be used with Somnivore because of its assisted machine learning design. We trained and tested the two applications on a reference data set that was selected from each of 6 bears, a one day recording in mid hibernation with a T_b_ at the peak of a multiday body temperature cycle [2], a one day recording in mid-hibernation near the trough of a body temperature cycle, and a one day recording in a non-hibernating state either in later summer or after full recovery from hibernation in early summer.

## Material and methods

### Reference data set

The analyzed data were collected for monitoring purposes during a previous study, and the procedures for animal care and implantation of the telemetry transmitters are described by Tøien et al. [2]. These experiments were approved by the Institutional Animal Care and Use Committee at University of Alaska Fairbanks (IACUC nos. 05-56 and 08-64). Sleep related data from this data collection have not been published previously. Briefly, American black bears (*Ursus americanos,* 62-93 kg) were captured in Alaska by Alaska Department of Fish and Game (ADF&G) during May–July, transferred to Fairbanks and held individually in an outdoor facility near the Institute of Arctic Biology in Fairbanks. The bears had been scheduled to be removed from the wild by ADF&G due to bear–human conflicts and would have been euthanized if not used by the project. In late August to late September in each year of the study, bears were immobilized with 8 mg/kg Telazol administered with a pole syringe and brought into surgery. Under isoflurane anesthesia and using aseptic protocol, the bears were implanted with telemetry devices (model T28F-14B, Konigsberg Instruments Inc., Pasadena, CA 91107) for measurement of T_b_, EKG, and deltoid muscle EMG. Using the same devices, global cortical EEG was recorded with a single differential pair of electrodes diagonally 1 cm from the skulls midline seam and 2 cm in front of and behind the bregma. The electrodes were stainless steel bone screws that penetrated into the extradural space and were insulated with acrylic cement. A single pair of EOG electrodes was implanted subcutaneously above the eyebrows. Some of the bears were also implanted with a blood pressure transducer in the caudal aorta. During recording, EEG, EOG and ECG channels were filtered with a 1-pole high pass (HP) filter at 0.1 Hz, and the EMG channel with 4-pole 20Hz HP filter. All of these channels were filtered with 3-pole low pass (LP) filters at 150Hz. Bears were also implanted with 2–3 intraperitoneal temperature data loggers for additional measurement of T_b_ (TidbitT temp model UTBI-001, 0.06 °C resolution, Onset Computer Corporation Inc, Bourne, MA 02532). Both types of devices were calibrated against a mercury thermometer traceable to NIST standards in a water bath to an accuracy of 0.1 °C. During analysis additional digital filtering was applied as found appropriate by the manual sleep scorers, typically a 0.5 Hz 3^rd^ order Butterworth HP filter for EEG and a 1 Hz first order Butterworth HP filter for EOG.

Feeding was gradually removed over a two week period from mid October, and once showing signs of entering hibernation, the instrumented bears were transferred to undisturbed outdoor enclosures located in isolated forest lands near the Institute of Arctic Biology. The bears were maintained in artificial dens (91 × 97 × 98 cm inside dimensions, 865 L, constructed from welded 2.5 cm thick HD-polyethylene and insulated with 5cm Styrofoam) that functioned as respirometry chambers when the breakaway doors were kept closed. Breathing was recorded with total body plethysmography using custom designed differential pressure transducers based on Honeywell SDX05 pressure sensors with one side connected to the den interiors, and the reference side connected to spatial filters consisting of branched ca. 2 m long PP90 catheters ending about 1.5 m apart in the immediate vicinity of the dens. The pressure signals were LP filtered at 0.1Hz and also HP filtered with a 1000 s time constant that prevented long term drift off the baseline. Radio telemetry signals were received and decoded by a base station (Konigsberg TD14), and recorded at 500 Hz with a data acquisition system (CA recorder, DISS LLC). The originally recorded CArecorder files (now an obsolete file format) were translated to European Data Format (EDF) using custom code written in open source Lazarus Free Pascal by the first author, adapting the open source PUMA repository (Dr. Johannes W. Dietrich, Ruhr University of Bochum) to write large EDF files. Adhering to EDF/EDF+ standard, the data were split into animal specific 1 day long files and relevant metadata for the signals added.

We selected from each of 6 bears three 24 h recordings: from mid-hibernation with a T_b_ at the peak of a multiday body temperature cycle [2] (T_b_= 34.9 ± 0.5°C), from mid-hibernation near the trough of a T_b_ cycle (T_b_= 32.1 ± 0.7°C), and from a non-hibernating state either in early or late summer (T_b_= 38.1 ± 0.4°C). These data were manually qualified by 3 different volunteer scorers. Scorers #1 and #3 had previous publication records in sleep research that included manual scoring of EEG data from mammals [20, 21, 22, 23]. Scorer #2 recorded all the data to be analyzed and had the longest experience with sleep data from hibernating bears, including the real-time observation of polysomnograms with video monitoring. None of these scorers were authors of the software that was used. As common in animal studies, we scored data into stages of NREM, REM, and Wake using 10 s epochs, and generally mapped what would be human stage N1, N2 and N3 to NREM, otherwise we used REM and Wake. We instructed scorers to not base scores on preconceptions about allowed transitions, i.e. not preventing Wake to REM transitions. In our preliminary analysis we also evaluated whether to also score a transition stage of Drowsiness with intermittent medium amplitude slow waves during inspirations occupying less than 50% of an epoch. However, we found difficulty in obtaining consistency among scorers, and it did not show consistent differences in and out of hibernation.

While we used the Polyman EDF reader as default viewer, we did not restrict use of other viewers or which additional channels that could be used for the manual scoring beyond EEG/EOG/EMG. Thus scorer #1 used Somnivore (with automatic scoring disabled) as viewer. Additional information from color spectral density arrays generated by the open source EDFbrowser (Teunis van Beelen, https://www.teuniz.net) was also used for assistance in the evaluation of the data. The scores were saved as separate EDF+ compliant files and were also exported in text format. From the qualifications of the 3 individual scorers, we calculated consensus scores though a voting system using code written in Lazarus Free Pascal. The single set of consensus scores were used in the further software training and testing.

### Machine learning training

Due to the differences in design of the software, training differed between Somnivore and Somnotate. For training Somnivore we used 100 epochs of NREM, REM, and Wake randomly selected by the software from each file to be tested as per the design of the software. After an initial assessment we used only the EEG and EOG channels for training as the recorded EMG from the deltoid muscle showed very little activity in bears during hibernation, possibly due to the typical curled postures of the bears [1] that causes stretching of the neck muscles. We applied a scoring rule of minimum consistency of 3 epochs for each of NREM, REM and Wake, which was necessary to approximate the sensitivity to changes in signal frequencies during transitions between states. Score Forcing that was an available option was not used to prevent Wake to REM transitions as it was found that spurious qualifications of short periods of Wake in the NREM to REM transition could affect what was correct detection of REM periods.

Somnotate was trained on multiple complete one day recordings using the EEG and EOG channels. We used a time resolution of 10s (same as the epoch length), which was found to be a good compromise to prevent too high sensitivity to state changes during transition states. For initial assessment we used multiple training data sets. In non-holdout mode we used all the available data in a training data set, while in holdout mode we trained Somnotate with all the files in the training data set except the specific file to be tested. We created training models: a) based on a training data set with all the reference data, b) separate training data sets for hibernating and non-hibernating states, and c) separate training data sets for the files at high and low T_b_ within the hibernating state. These were tested for overall accuracies in Somnotate.

### Statistical analysis

The metric used for detailed evaluation of algorithm generalization accuracy versus manual scoring for both applications was F-measures calculated after import of scoring results into Somnivore. It is one of the most stringent metrics available, as both high sensitivity and high precision are required for high F-measure [19, 24]. Data are given as means ± standard deviation. Further statistical analysis was performed in Sigmaplot ver. 11 and Excel. For testing of statistical significance between different tests on the same sets of animals we used two-tailed paired t-tests, otherwise two-tailed equal variance or unequal variance t-tests as found appropriate after testing the data distributions. Differences were considered significant at p<0.05.

## Results

Fig 1 shows examples with short segments of raw recordings of each vigilance state including a pronounced delta wave EEG pattern during NREM, low amplitude EEG with alpha waves and EOG with bipolar deflections during REM sleep, and quiet Wake including irregular low amplitude EEG with additional muscle activity during active Wake. When qualifying the transitions from Wake to NREM manual scorers followed the rule of 50% presence of medium to high amplitude slow wave; the example in Fig 2 shows an occasional period of episodic slow waves during inspiration and simultaneous shivering bursts occurring before NREM could be defined. We initially evaluated if we should score these periods as a Drowsiness stage, however an assessment by scorer #1 and #2 on samples from 1-2 bears revealed that there was too much uncertainty in defining the Drowsiness stage. Also, the relative occupancy was low and did not systematically change between hibernation and summer. Table 1 shows overall agreement of individual scorer’s qualifications with the consensus scores. Two scorers had overall agreement of 96.5% with the consensus, whereas the third scorer was lower at 90.5%. These differences reflected that scorer #3 had been able to spend the least amount of time to become familiar with the specifics of the data before scoring started. Only 0.5% of the total epochs were no-consensus epochs and these were typically in the transitions.

**Fig 1.**
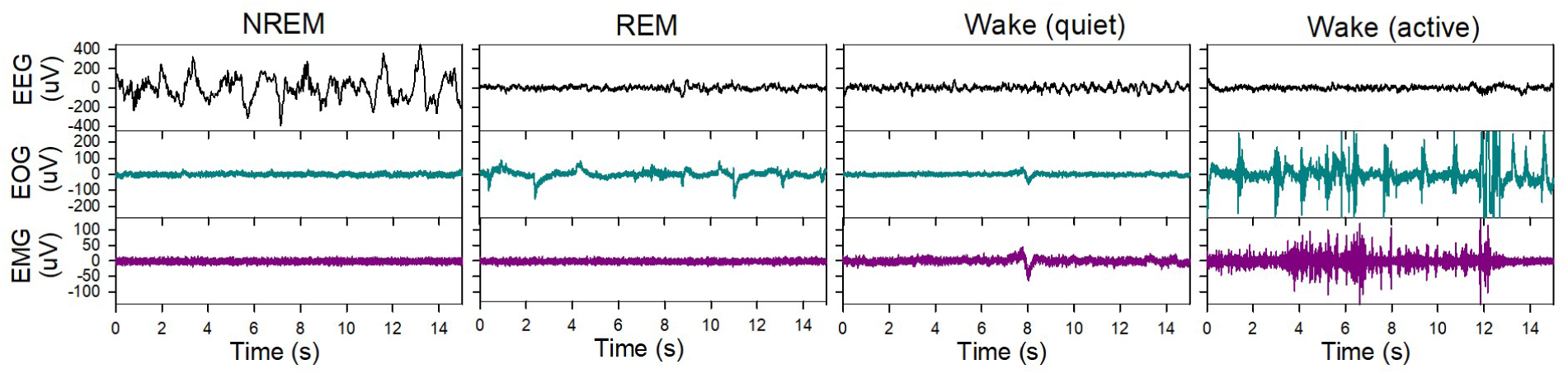
Short examples of raw recordings. These 15 s examples show EEG (top panels), EOG (middle panels) and EMG (bottom panels) during NREM, REM and different forms of Wake in a hibernating black bear.

**Fig 2.**
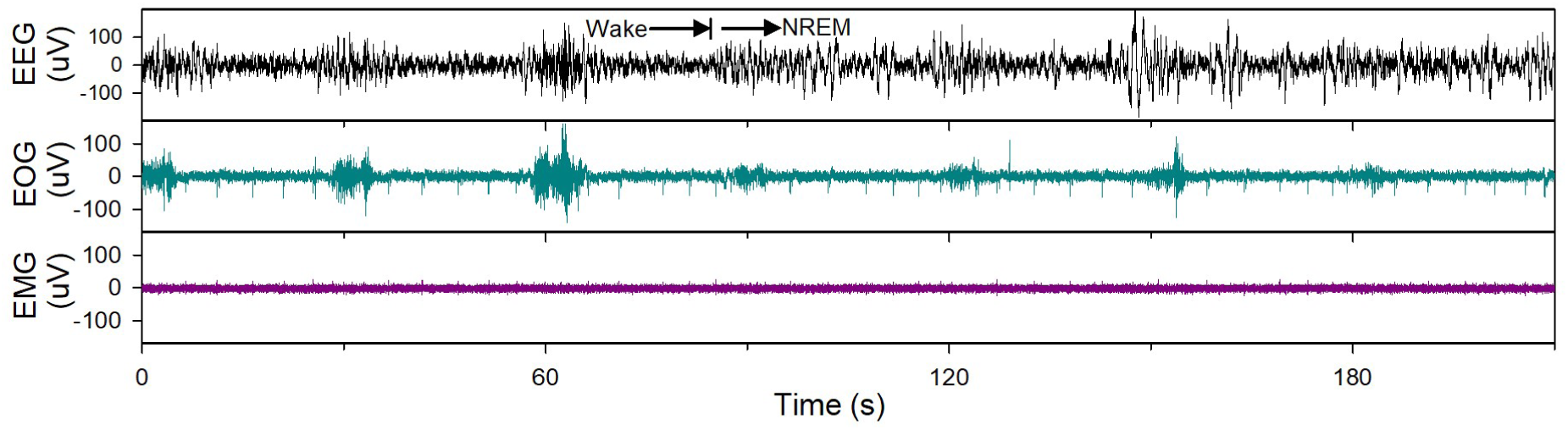
Wake to NREM transition. This raw recording example with EEG (top panel), EOG (middle panel) and EMG (bottom panel) shows a challenging transition from Wake to NREM in a hibernating black bear with discontinuous slow wave activity occurring during the inspiratory phase of breathing and that was initially too intermittent to be manually qualified as NREM. EMG activity from shivering bursts bleed into the EOG channel during breaths, while the EMG of the deltoid muscle was typically without activity.

**Table 1.** Agreement of individual manual scorers with the consensus.

|  | Total | Scorer 1 | Scorer 2 | Scorer 3 | Undefined | Average |
| --- | --- | --- | --- | --- | --- | --- |
| **Consensus epochs** | 154919 | 148778 | 148697 | 139447 | 776 |  |
| **Percent agreement** | 99.5% | 96.5% | 96.5% | 90.5% | 0.5% | 94.5% |

An initial assessment of how using different sets of training channels affected the overall accuracy of Somnivore is shown in Fig 3. There was a significant increase in accuracy of the hibernation sample from 91.4 ± 2.2% with training on the EEG channel only to 92.7 ± 1.6 % with training on both the EEG and EOG channels (p<0.05, paired t-test), while there was only a trend for increase in accuracy with more channels being used for training in summer samples (p>0.1). Thus, for further analysis, we standardized on using the EEG and EOG channels for training. While training Somnivore is normally performed on a subset of data from the file being tested, we also tested the additional capability to apply training data generated with other files, as shown in Fig 4. Accuracy significantly decreased from 92.7± 1.6 % with in-file training to 69.7 ±15.9% (p<0.001) when training on data from different animals during hibernation. Training on data from the same animal but from a file from a different T_b_ during hibernation decreased accuracy to 85.3 ± 5.6% (p<0.001). In summer bears accuracy decreased from 93.7 ± 3.2% to 84.9 ± 5.4 % (p<0.02) when training on data from a different animal than the one being analyzed. Thus, for further analysis with Somnivore, we only use training data from a subset of the file being tested.

**Fig 3.**
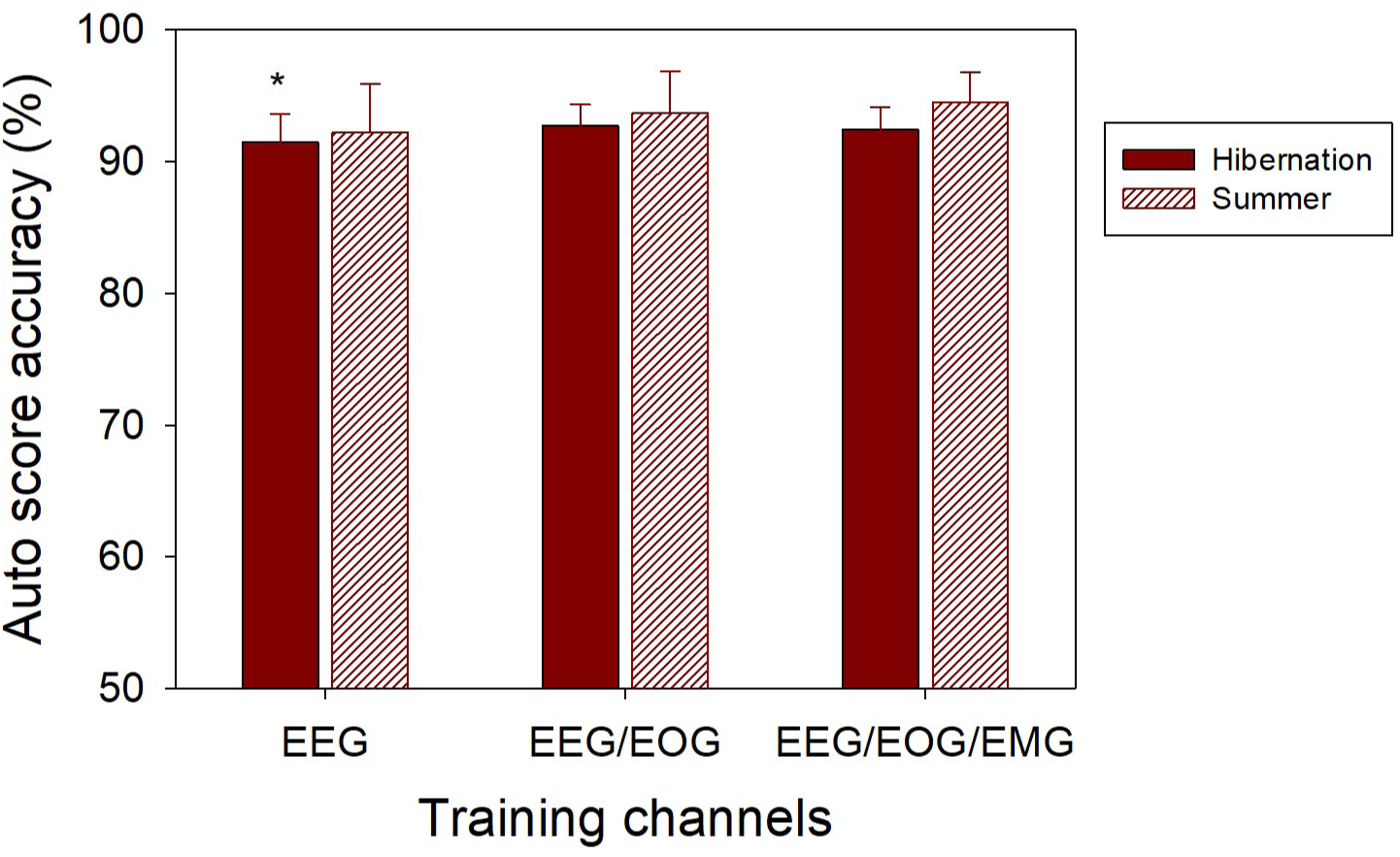
Somnivore overall scoring accuracy with different sets of training channels. Somnivore overall scoring accuracy when training on different channels of the analyzed file using a random selection of 100 epochs of each qualified state, showing different results for hibernation (solid bars) and summer samples (cross-hatched bars). * designates a significance level of p<0.05 compared to using EEG/EOG channel for training.

**Fig 4.**
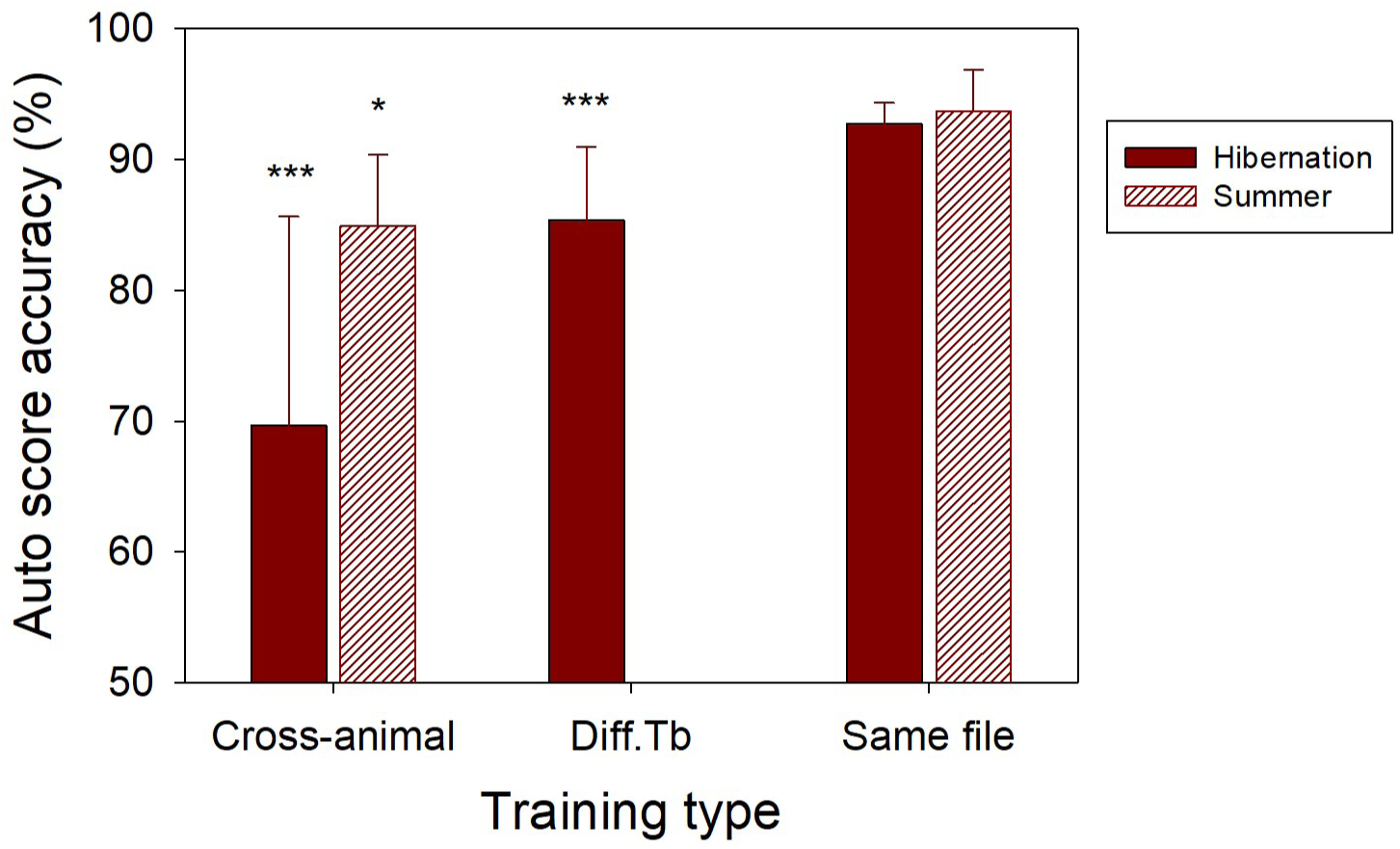
Somnivore overall scoring accuracy when training on different files. These tests used the EEG and EOG channels for training, but training was performed on a randomly selected file from a different animal in the reference data set (Cross-animal), or a different file from the same animal at different T_b_ level (Diff. Tb), or the standard training on data from the file being analyzed (Same file). Solid bars designate analyzed and training data from hibernation, cross-hatched bars designate analyzed and training data from summer. Stars designate significant differences compared to standard training on the analyzed file, * p<0.05, *** p<0.001.

The ability of Somnotate to train on multiple files and apply the training model to files recorded under different conditions and in particular files recorded at different T_b_ called for additional tests of the robustness of the training data. Fig 5 shows the results of testing with different training models, both with all files in the data set included and with holding out the tested file from the training data. Holdout testing only decreased overall accuracy of Somnotate by 1.1-1.3% (p<0.05 to p<0.0001) except 4.7 % decrease in holdout accuracy when training on both hibernation summer files (p<0.05). Best accuracy for hibernation data was obtained by training on hibernating files only, 92.0 % for holdout training and 93.2 % for non-holdout training. Training separately on hibernation files with high T_b_ (from the peak of the multiday body temperature cycles) and low T_b_ (from the trough of the body temperature cycles) did not improve overall accuracy (p>0.37). Summer data had significantly lower accuracy when trained on both hibernation and summer files (82.7% for holdout testing and 87.4% for non-holdout testing) compared to training on summer data only (93.6% for holdout testing and 94.9% for non-holdout testing, p<0.05).

**Fig 5.**
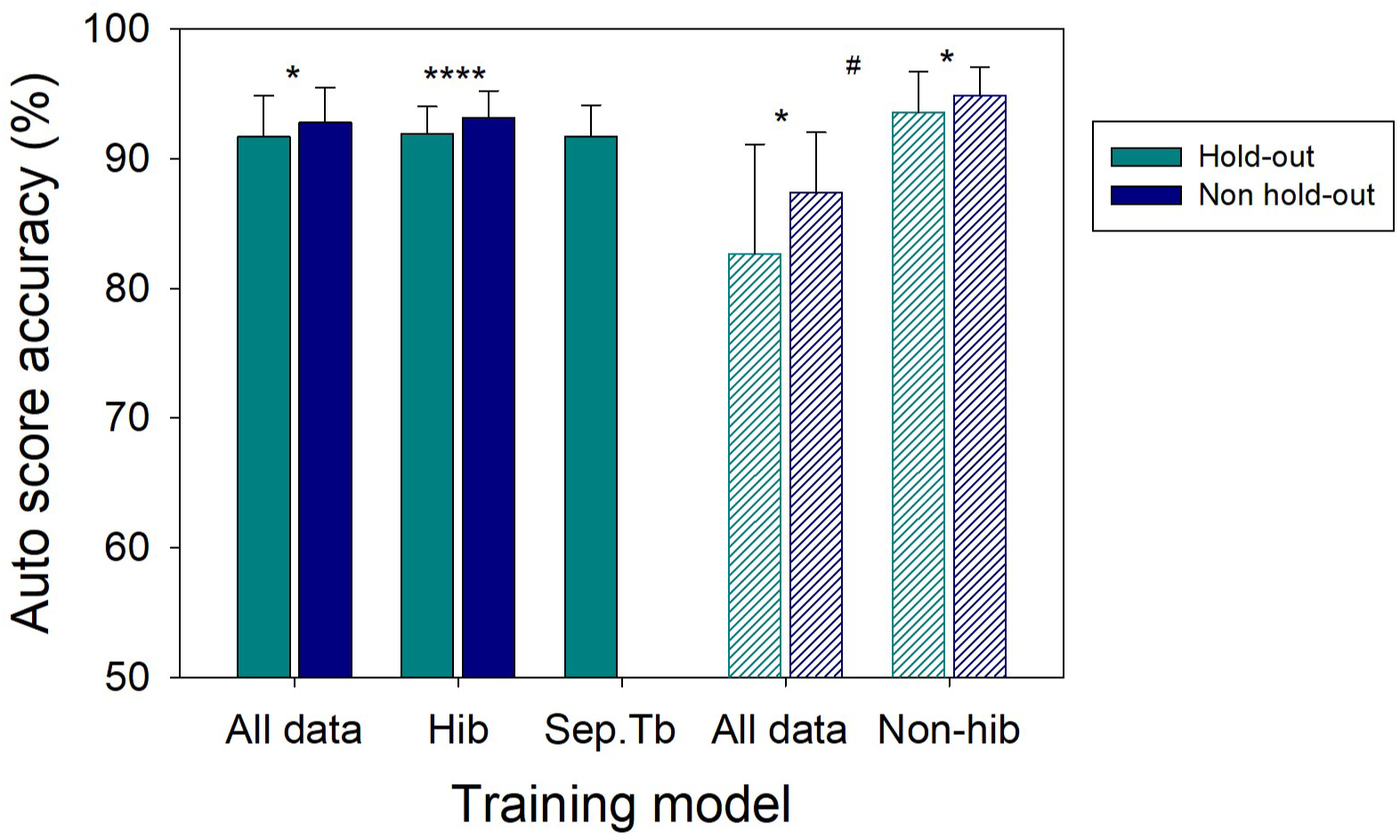
Somnotate overall scoring accuracy when training on different data sets. Color designates if tested file is held out from training data set (hold-out training, teal color) or not held out from training data set (non hold-out training, blue color). Bar pattern designates if the tested files is from hibernation (solid bar) or from summer (cross-hatched bars). Bar labels designate if both hibernation and summer samples were used for training (All data), or hibernation samples were used for training (Hib), or if training was performed separately on hibernation samples from high and low T_b_ (Sep Tb), or summer samples were used for training (Non-hib). Significant differences between hold-out and non-holdout training is designated by * p<0.05, **** p<0.0001. Significant differences between training on all data and summer samples only (both hold-out and non-holdout) is designated by # p<0.05. The EEG and EOG channels were used for all training.

Further detailed testing of both applications was performed without holdout of the tested file from the training data. Fig 6 shows examples of one-day time courses of manual consensus scores alignment with the automated scores from Somnotate and Somnivore and color density spectral array plots visualizing changes in frequency distributions of the EEG signals. Good alignment is seen between automated scores from both applications with consensus scores both in the summer file and hibernation file, with a few deviations in the transitions between states. The detail from hibernation exemplifies a troublesome NREM to REM transition where the manual scorers did not reach a consensus and the automated scores inserted some wake at the beginning and into the REM period. Analysis of transitions of the full data set indicated a much higher prevalence of Wake to REM transitions with Somnivore (0.33) than with Somnotate (0.15) and both were considerably higher than the manual consensus scores (0.03). However, the duration of these Wake before REM periods were usually very short in the 0-5min bin and thus would not be expected to contribute much to the total stage occupancy.

**Fig 6.**
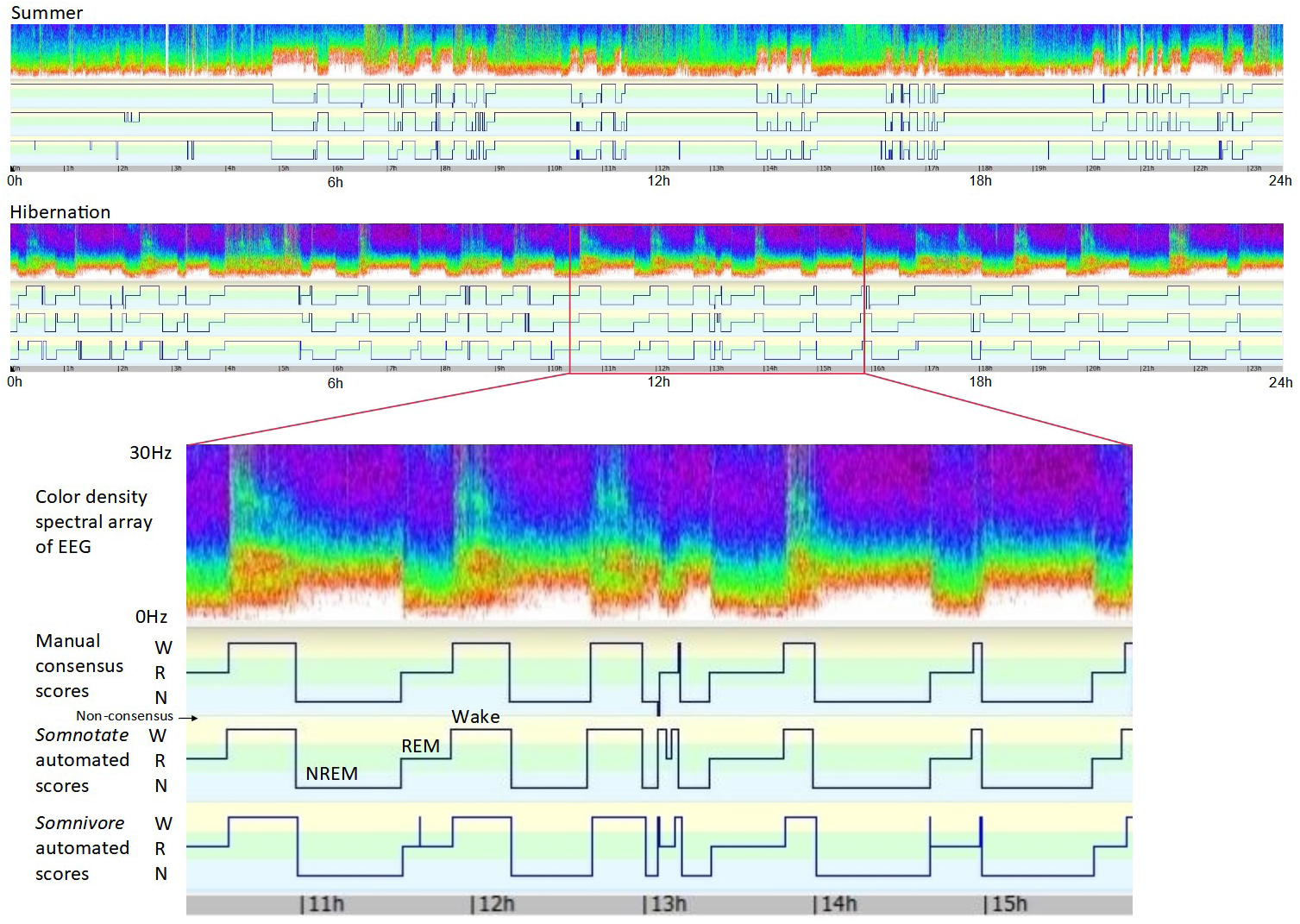
Examples of manual consensus scores vs. automated sleep scores from 24 h periods in the same bear in summer and hibernation. For each, top panel: color spectral density plot with frequency (0-30Hz) on y-axis and non-calibrated intensity color coded; middle panel: consensus manual sleep scores; bottom panel: automated scores by Somnotate using separate training data for hibernating and non-hibernating bears. Each trace shows Wake (W) as highest value, REM (R) in the middle, NREM (N) as low value and non-consensus scores as low off-scale deflections. The 5.5 hour detail from hibernation includes an NREM to REM transition that proved problematic for both manual scorers and the automated scoring programs.

A summary of F-measures for comparison of automated scores with the consensus scores for the different vigilance states is shown in Fig 7. There were no significant differences in F-measures between Somnivore and Somnotate (p>0.18). Both had overall average F-measures in the 0.90-0.95 range. NREM sleep had the lowest variability of the F-measures, indicating a tendency for the automated scoring to have less sensitivity to variability in the data for this state. REM sleep detection in summer data had the lowest average F-measure of 0.84± 0.12 with Somnivore.

**Fig 7.**
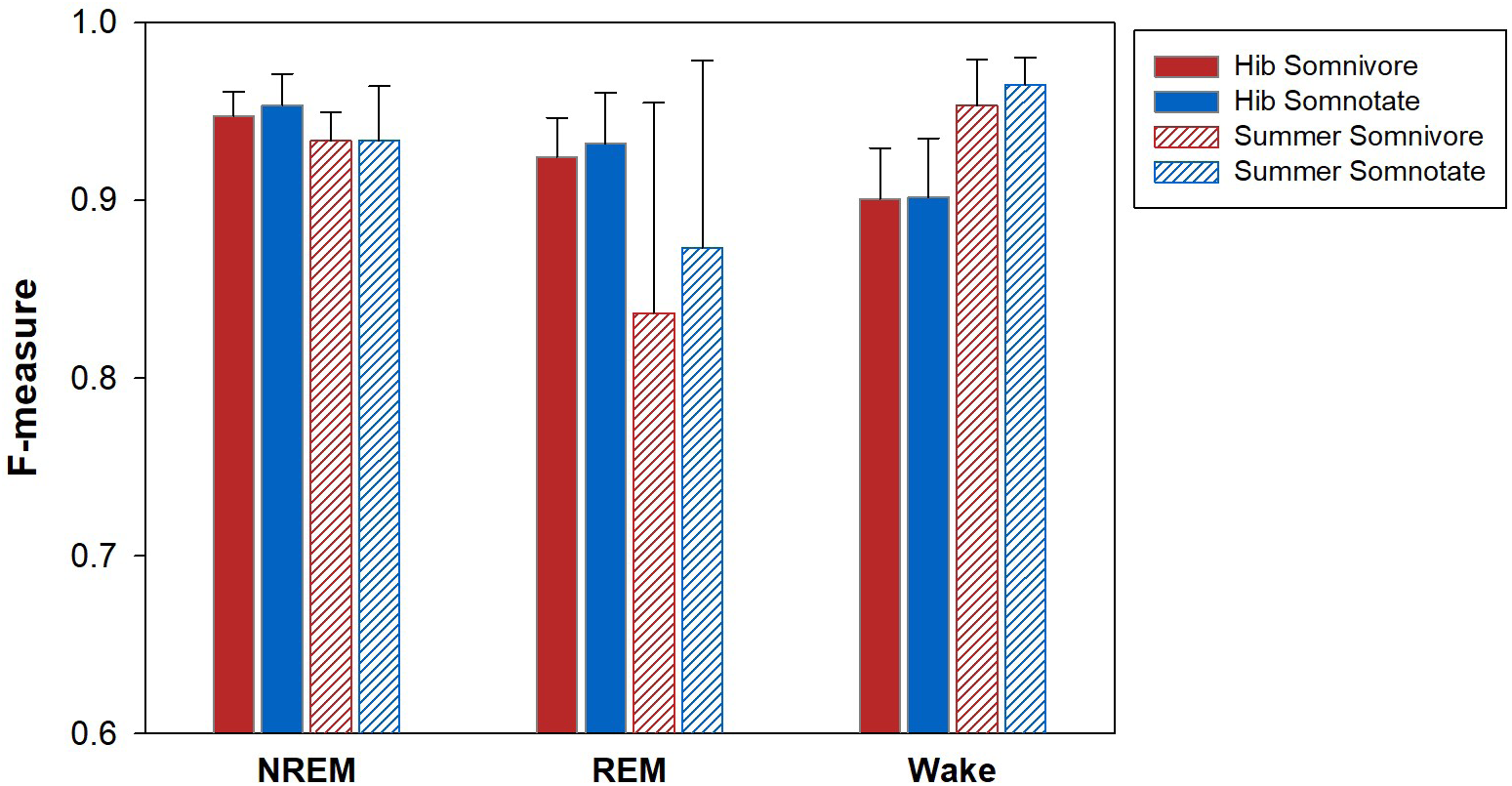
F-measures of automated scoring vs consensus scores for different vigilance states. F-measures of NREM, REM and Wake for Somnivore (red bars), Somnotate (blue bars), in hibernation (solid bars) and summer (hatched bars), positive error bars indicating SD.

Fig 8 shows the average distribution of vigilance states in hibernation and summer for the reference data set from 6 bears based on manual consensus scores and automated Somnivore and Somnotate scores. Bears spent 1.95x as much time in NREM, 2.26x in REM, 2.05x in total sleep, and 0.51x in Wake during hibernation compared to summer based on the consensus scores. The distribution of vigilance states in hibernation based on the consensus scores was NREM 42.9 ± 7.0 %, REM 22.0 ± 2.3 %, total sleep 64.9 ± 6.7 %, and Wake 34.6 ± 6.7%. The automated scores in hibernation did not differ significantly from the consensus scores (p> 0.05 to p>0.52). During summer, the distribution of vigilance states based on consensus was NREM 21.9 ± 3.6 %, REM 9.7 ± 2.8 %, total sleep 31.7 ± 4.9 %, and Wake 68.0 ± 4.8 %. All vigilance states differed significantly between hibernation and summer for both consensus scores and automated scores (equal variance t-test, p< 0.0001, n=12 and 6 respectively). There were no significant differences in detected vigilance states between the two automatic scoring methods (p=0.08-0.97, paired test). Both applications consistently overestimated sleep times in summer by a small amount (NREM 2.4% for both, REM 3.3% for Somnivore and 1.9% for Somnotate) and underestimated wake times by a small amount (5.3% and 3.9%, respectively) compared to the consensus scores (p<0.05 except p=0.13 for REM with Somnotate, paired t-tests). Small overestimates of sleep times remained even when including EMG among the training channels for Somnivore (p<0.05).

**Fig 8.**
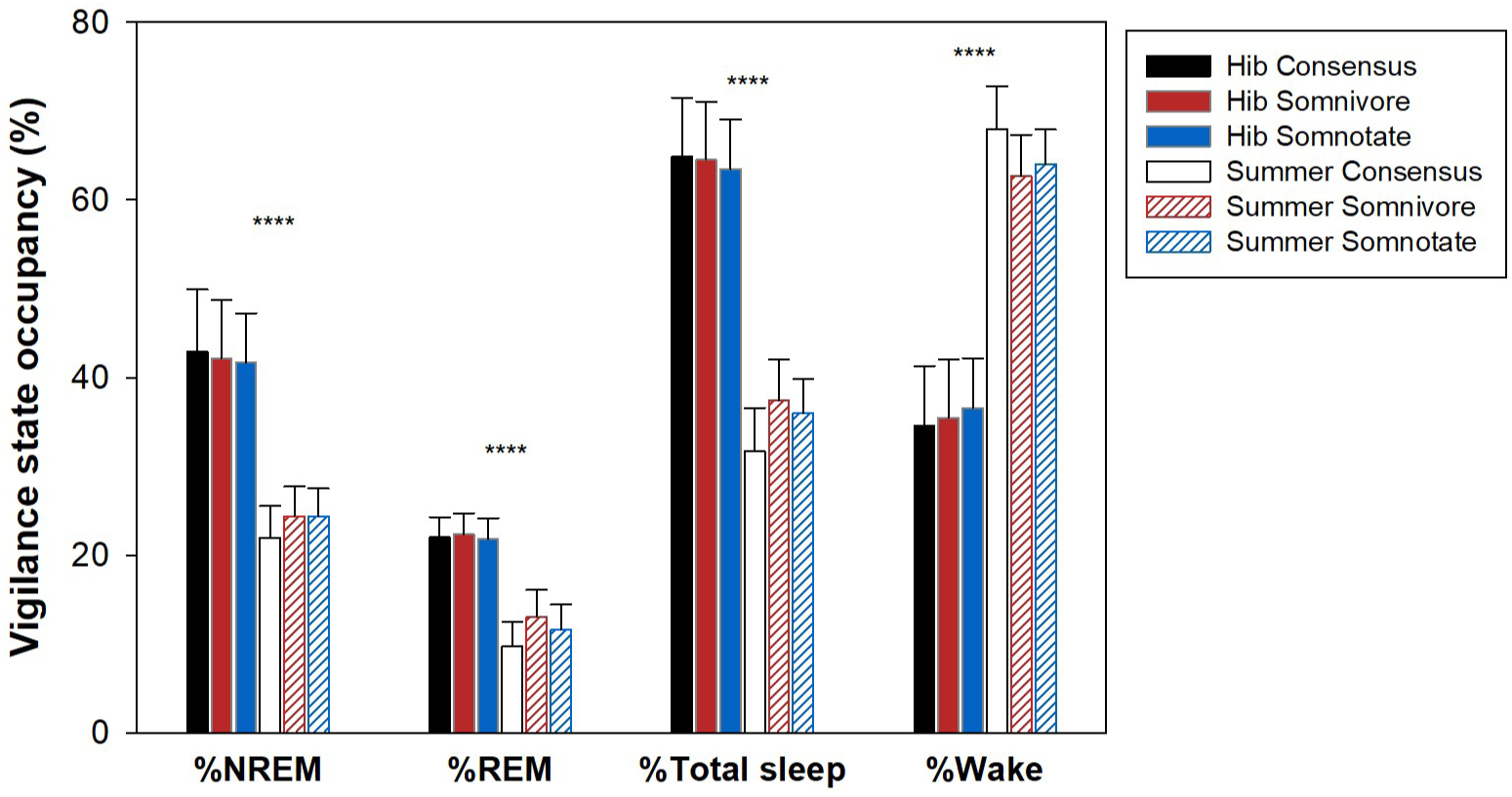
Distribution of vigilance states in the reference data set from 6 bears in hibernation and summer with different scoring methods. Colors show manual consensus scores (black or white bars), automated Somnivore scores (red bars), Somnotate scores (blue bars). Bar pattern designates hibernation (solid bars) and summer (open or hatched bars). Positive error bars indicate SD. Significance level for comparison of hibernation to summer is indicated by **** at p< 0.0001 and applies to all three data sets.

A more detailed distribution of the performance of the two applications is visualized in Fig 9 showing individual files F-measures of Somnivore vs. those of Somnotate. F-measures with both applications are clustered in the 0.90-0.98 range. Outliers with low F-measures were consistently low for the same files with both scoring applications; correlations of F-measures between Somnivore and Somnotate were for NREM R^2^ = 0.73, REM R^2^ = 0.57, Wake R^2^ = 0.78. For Wake and NREM, the outliers were mostly distributed among the hibernation data (closed symbols), while outliers for REM were for summer data (open symbols).

**Fig 9.**
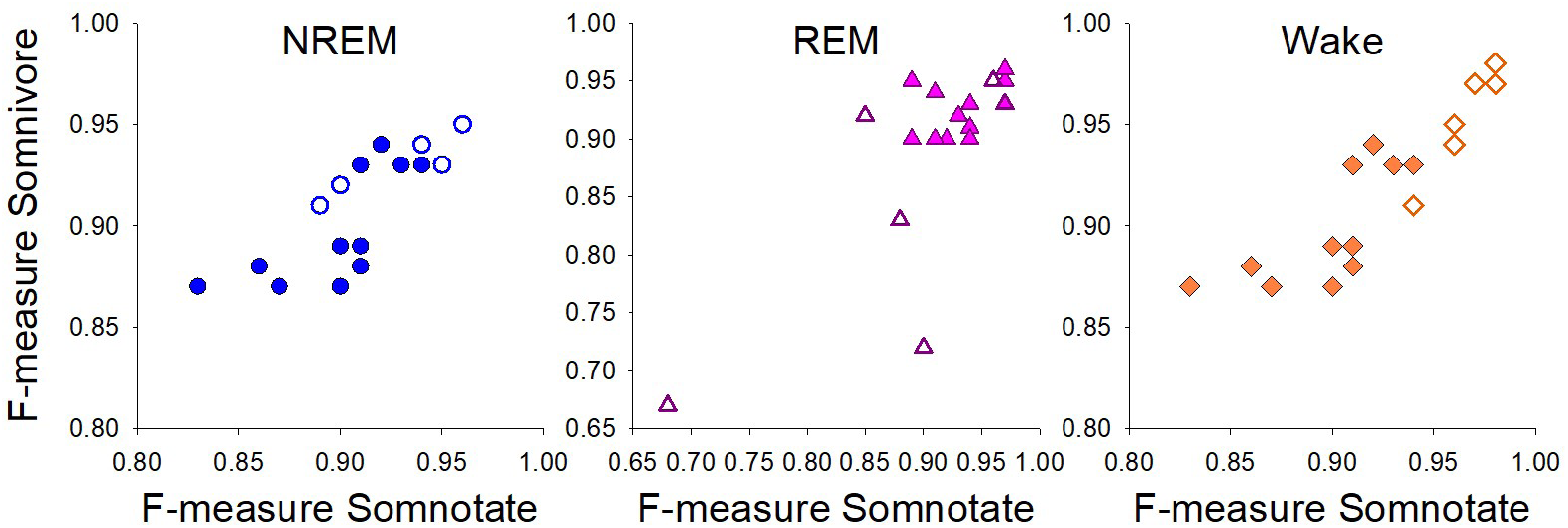
Detailed F-measures of Somnivore vs Somnotate for different vigilance states. Hibernation (closed symbols), summer (open symbols). The F-measures of automatic scores vs consensus scores show that outliers with low F-measures tended to occur with both Somnivore and Somnotate on the same files.

## Discussion

Comparing the use of two different automated scoring applications, this study is the first to score vigilance state recordings from a non-human large animal model that includes hibernation and non-hibernation. A major concern with the data from hibernators is the frequency shifts in the EEG signal with decreases in brain temperature [17] that could cause problems for correct designations of vigilance states by the automated scoring algorithms. The drop in T_b_ of hibernating black bears is more moderate, less than 8°C [1], than it is in smaller hibernators where it can exceed 30°C. It is not known if those changes as well as the multi-day body temperature cycles during the hibernation season [2] could prevent correct detection of vigilance state by the machine learning algorithms. In addition, it was not known if the nature of sleep during hibernation differs in other qualities from that during summer in ways that could compromise integrity of the automated scoring. Our results show that with proper use of training models the automated scoring with the two tested machine learning based qualifiers show remarkably similar performance and can approximate the performance of manual scorers.

The use of consensus between 3 manual scorers in the present study helped reduce uncertainties due to subjective judgment of the manual reference scores. As shown by Brodersen et al. [18] a significant increase in accuracy of automatic scores was obtained already when increasing the number of manual scorers from 1 to 3. As applied, the assisted machine learning model of Somnivore [19] was unproblematic with respect to variability in the data as it had the advantage of training on data very similar to that being analyzed by using a limited number of epochs from each file to be analyzed as training data. Our protocol of using 100 epochs per vigilance state, which is above the number that was found to provide optimal performance on human data and near optimal in rodents [19], only requires manual scoring of 3.5% of the total data to be automatically scored. Our initial testing showed relatively little dependence of accuracy on which training channels were used. While it was optimal to train on EEG and EOG channels in hibernation, it appears that the most important information during training is extracted from the EEG channel. Thus, only a 1.3% decrease in overall accuracy is predicted if an EOG channel is not available in the recording (Fig 3). It was not surprising that training Somnivore on data recorded from a different bear than the analyzed file (cross-animal training, Fig 4) caused a profound decrease in overall accuracy for the hibernation data, as it included both the effects of a different animal and T_b_ and is beyond the design of Somnivore. Training Somnivore on files from the same animal at a different T_b_ during hibernation also caused a marked decrease in overall accuracy. Thus, for hibernating animals Somnivore is best used with assisted machine learning as designed.

After training Somnotate on 6 one-day recordings of mouse data, it was shown to be highly robust and tolerated as much as 50% randomly mislabeled epochs added to the training data before accuracy decreased [18]. For our bear data we found that overall accuracy was decreased if both hibernation and summer data were used for training and applied to summer data, while we did not find a substantial drop in accuracy when the same training was applied to hibernation data (Fig 5). This may partly reflect that our reference data set consisted of two times as much data from hibernating bears as non-hibernating bears, but could also be due to other characteristics of the signals that differ too much between hibernation and summer. The finding of a high accuracy on hibernation data with training models based on both data from high and low T_b_ during hibernation is consistent with the previously observed robustness of Somnotate on mice data [18]. The ability to use a single training model within hibernation in bears is an important finding, as it otherwise would have been difficult to define which training model should be selected when T_b_ is varying though a continuum during multi-day body temperature cycles.

The overall accuracy of automatic scores of Somnotate with separate training models for hibernation and summer was only slightly lower or at the level of agreement of single manual scores with the manual consensus scores (92.0-93.5% vs 94.5% Fig 5 and Table 1, respectively). The use of consensus scores for training likely helps avoid outliers in the training data due to subjective judgment and carries over to improve the automated scores. Our results compare favorably with published results on machine learning based automated sleep scoring that had overall accuracies in the 80-98% range for human data [13, 14, 15, 16]. The detailed F-measures also indicate excellent ability to distinguish between vigilance states for both for Somnotate and Somnivore with F-measures mostly in the 0.90-0.98 range. The two applications performed remarkably similarly with a tendency for lower average F-measures and more spread in F-measures for REM in summer (Fig 7). The lower occupancy of REM in summer (Fig. 8) may have inherently influenced the F-measures. Previous testing of Somnotate on mouse data resulted in average F-measures of 0.97, while tests of two other applications on the same mouse data set had F-measures of 0.94 and 0.73 respectively [18]. Somnivore tested on rodent data had highest average F-measures for Wake and NREM at 0.95 and 0.94, respectively, with REM lowest at 0.91 [19]. This is similar to the lowest average F-measure of about 0.90 for REM with both Somnotate and Somnivore in our study, while NREM had the highest average F-measure of 0.94-0.95 (Fig. 7).

The total distribution of vigilance states based on automated scores of the reference data set show for the first time evidence that bears sleep about 2x as long during hibernation compared to summer (Fig 8). The functional physiological base for this increase in NREM and REM compared to summer will require a much deeper analysis than offered by our limited analysis here. However it is unlikely that bears share functional similarities with the increase in delta waves reported during early arousals in smaller hibernators [9,10], where the resolution of that SWA is resolved without sleep [11, 12, 25]. This elevated SWA after arousal from low T_b_ in smaller hibernators may be related to synaptic loss during deep torpor that occurs linearly with a decrease in brain temperature and could be due to a decreased release or responsiveness to wake-promoting factors or to the slow regrowth of exitatory synapses in the cortex and thalamus [26]. These sleep changes observed in deep hibernators have limited translational relevance to hypometabolic states at brain temperatures closer to normal as in bears (30-35°C) [2] where T_b_ is high enough to allow cortical brain activity to be recorded continuously and loss of synaptic connectivity is unlikely to occur.

The results are a snapshot of two days during the coldest period during mid hibernation when sleep patterns could have been affected by shivering. Thus, scoring extended periods during hibernation in more bears is needed to better understand the time courses of sleep times and bout lengths in relation ambient temperature and the multiday body temperature cycles [2]. These results are remarkably similar for Somnivore and Somnotate and did not differ significantly from the manual consensus scores during hibernation (Fig 8). However, there was a small but significant overestimate of sleep time and underestimate of wake for the summer data compared to the manual consensus scores. The summer data might present some challenges in that the patterns were more fragmented, with bears drifting in an out of drowsiness and sleep. This could perhaps cause a cumulative effect if automatically scored sleep periods extend one or a few more epochs during each transition than they do in the manually scored recordings. The summer recordings were also less clean with presence of artifacts as bears moved around including EMG infusion into the EOG and EEG recordings. Also, there were more problems with telemetry signal dropouts in summer recordings, although both programs have strategies to detect and exclude these epochs.

Problematic files with lower F-measures tended to be outliers in both Somnivore and Somnotate (Fig 9). It is likely that some data files have less well-defined states than others making the automatic scoring more difficult. An alternative or additional explanation is that these problematic files were also difficult for the manual scorers resulting in lower quality manual consensus scores, and that the automatic scoring picked up features not recognized by the manual scorers, resulting in the automatic scores deviating from the manual scores.

A particular potential difficulty is the correct detection of transition states. As pointed out, the entry into NREM in hibernating bears could be challenging to qualify due to the respiration-related periodic appearance of slow waves (example Fig 2). However, it did not seem to affect reliable detection as NREM had the highest F-measures (Fig 7). Scoring of transition states of NREM was generally similar between the automatic scores and the consensus manual scores (examples in Fig 6), and misalignment of a few epochs would have minimal effect on the overall distribution of vigilance states except for potential cumulative effects caused by fragmented patterns with frequent transitions. Automatic detection occasionally failed to detect correct transitions as determined by the manual scorers, particularly with respect to transitions into REM. This was particularly the case with Somnivore. However the incorrectly detected periods of Wake before REM were quite short and likely did not contribute much to overall occupancy of vigilance states. The sensitivity of the scoring software to state changes caused by short term frequency changes in the EEG is modulated by the configuration. For Somnotate we used a sampling interval of 10 s, and for Somnivore a scoring rule of minimum score consistency of 3 epochs was required to allow a state change for all three states. These are settings used to align scoring rules to the conventions used to manually score the data, but do not necessarily reflect the short term underlying frequency changes. They might compensate for uncertainty in the determinations that otherwise would cause flipping of states back and forth and not considered state changes by manual scorers. However if the required minimum score consistency in Somnivore were set too high or the defined sampling interval in Somnotate was set too long, the responsiveness to state changes could become too slow. So definition of these parameters are compromises, but can possibly be fined tuned. For the manual scores we decided to not base scores on pre-conceived conventions about allowed state changes, i. e. not excluding Wake to REM transitions as making a priori assumptions may not be well founded when scoring a new species and in a hibernating state. We did detect some unusual Wake to REM transitions in the bears, but detailed anaysis of more data would be needed to draw any conclusions on validity and origins. Caution should be used and not just reliance on automatic scores to make conclusions about presence of unusual transitions as exemplified in Fig 6. Somnotate has the additional ability, not explored in the results presented here, to flag epochs with low probability for the state determination [18] that could be useful in finding transition states that need manual inspection.

To conclude we have for the first time shown that automatic sleep scoring of a large mammal comparing hibernation and non-hibernation states can be done with trained machine learning models. Both Somnotate and Somnivore have overall accuracies approaching the best in the literature based on automatic vs. manual consensus scores. We now have the tools to determine distribution of vigilance states in the data set of 3500 days of polysomnographic recording from hibernating and non-hibernating black bears using automated batch scoring. We think one should keep an open mind on the possibility of unusual vigilance state transitions in a new species and in unexplored states, but we suggest that conclusions about problematic transitions should be determined by manual inspection that can be assisted by software support.

## Acknowledgements

We thank Stephen Morairty for early comments on the data and the manual sleep scoring analysis. Alaska Department of Fish and Game helped providing bears as previously described [2].

## Data availability

The data underlying the results presented in the study are available from the National Sleep Research Resource (https://doi.org/10.25822/cp4n-fh44) at https://sleepdata.org/datasets/toien-2025. The code for Somnotate is available at https://github.com/paulbrodersen/somnotate. The alaska branch contains custom modifications for the project. The Somnivore code is proprietary. The code used for extracting consensus scores is available at https://github.com/otoien/ConsensusCode.

